# Distribution and frequency of salivary gland tumours: an international multicenter study

**DOI:** 10.1101/2022.03.17.484728

**Authors:** Ibrahim Alsanie, Shahad Rajab, Hannah Cottom, Oluyori Adegun, Reshma Agarwal, Amrita Jay, Laura Graham, Jacqueline James, A William Barrett, Willie van Heerden, Mariano de Vito, Alessandra Canesso, Akinyele Olumuyiwa Adisa, Abdul-Warith Olaitan Akinshipo, Oluseyi Folake Ajayi, Mark Chukwuemeka Nwoga, Chukwubuzor Udokwu Okwuosa, Olufemi Gbenga Omitola, Efetobo Victor Orikpete, Merva Soluk-Tekkesin, Ibrahim Bello, Ahmed Qannam, Wilfredo Gonzalez, Maria Eduard Perez de Oliveira, Alan Roger Santos-Silva, Pablo Agustin Vargas, Eu-Wing Toh, Syed Ali Khurram

## Abstract

**Background:** Salivary gland tumours (SGT) are a relatively rare group of neoplasms with a wide range of histopathological appearance and clinical features. To date, most of the epidemiological studies on salivary gland tumours are limited for a variety of reason including being out of date, extrapolated from either a single centre or country studies, or investigating either major or minor glands only.

**Methods:** This study aimed to mitigate these shortcomings by analysing epidemiological data including demographic, anatomical location and histological diagnoses of SGT from multiple centres across the world. The analysed data included age, gender, location and histological diagnosis from fifteen centres covering the majority of the world health organisation (WHO) geographical regions between 2006 and 2019.

**Results:** A total of 5798 cases were analysed including 65% benign and 35% malignant tumours. A slight female predilection (54%) and peak incidence between the fourth and seventh decade for both benign and malignant tumours was observed. The majority (69%) of the SGT presented in major and 31% in the minor glands. The parotid gland was the most common location (70%) for benign and minor glands (46%) for malignant tumours. Pleomorphic adenoma (70%), and Warthin’s tumour (17%), were the most common benign tumours whereas mucoepidermoid carcinoma (25%) and adenoid cystic carcinoma (16%) were the most frequent malignant tumours.

**Conclusions:** This multicentre investigation presents the largest cohort study to date analysing salivary gland tumour data from tertiary centres scattered across the globe. These findings should serve as a baseline for future studies evaluating the epidemiological landscape of these tumours.

## Introduction

Salivary gland tumours (SGT) are a heterogeneous group of neoplasms with a wide range of histological subtypes making diagnosis challenging for pathologists. Fortunately, they are rare, with an annual estimated incidence of approximately 2.5–3.0 per 100 000 people in the Western world [1].

Most SGT are benign with ~70% arising in major glands and ~25% are from the minor glands. Malignant SGT comprise approximately 2-6% of all head and neck cancers [1,2] with 15-35% of parotid gland, 41-45% of submandibular and 70-90% of sublingual glands tumour being malignant [3]. In comparison, more than half of **t**he minor glands tumours (including palate, tongue, the floor of the mouth, retromolar region and lips) are likely to be malignant [3,4,5]. Other rare sites for SGT include the larynx, trachea, lacrimal glands, nasal cavity and heterotopic salivary tissue within the mandible and the lymph nodes [4]. Tumours involving minor glands have worse prognosis, higher recurrence rate and poor outcomes compared to major gland tumours [6].

To date, numerous studies have reported epidemiological data for SGT. However, they are somewhat out of date or reflect relatively small datasets from a single centre or local population only [4–5,7–16]. One of the most recent studies from 2012 reported incidence of these tumours in two distant geographical locations without detailed comparative analysis [17]. These shortcomings necessities the need for epidemiological evaluation of SGT from multiple centres preferably different geographical locations across the world with a view to analysing the distribution of different subtypes of SGT as well as identifying trends in the different populations.

Therefore, the aim of this multicentre international study was to analyse SGT data from numerous tertiary hospitals across the world with a view to obtaining up to date frequency and distribution of SGT. Further investigation of demographic and anatomical location of SGT and correlation of findings from different geographical locations was also performed.

## Material and methods

All salivary gland tumours diagnoses between 2006 to 2019 were retrieved from the Pathology databases of the involved Oral and Maxillofacial/Head and Neck Pathology departments. The year 2006 was selected in this study as cut-off year for two main reasons; 1) A number of SGT overviews were published prior to this including analysis of a large cohort from the Lead Institute and 2) Since then there have been significant changes to histological classification of SGT. The UK centres included Sheffield (School of Clinical Dentistry, University of Sheffield and Sheffield Teaching Hospitals NHS Foundation Trust), East Grinstead (Queen Victoria Hospital), Belfast (Belfast Health and Social Care Trust/Queen’s University Belfast), and two centres in London (Royal London Hospital and University College London Hospital). The international collaborators included centres from Italy (Department of Pathology, Camposampiero), Turkey (Department of Tumour Pathology, Istanbul University), Nigeria (including Ibadan, Lagos, Enugu and Port-Harcourt), South Africa (University of Pretoria), Saudi Arabia (Department of Oral Medicine and Diagnostic Sciences, College of Dentistry, King Saud University, Riyadh), Brazil (Piracicaba Dental School, University of Campinas, São Paulo State) and Chile (Universidad de Valparaíso, School of Dentistry) (Figure 1). These were split into regions for some of the comparative analysis.

**Figure 1:**
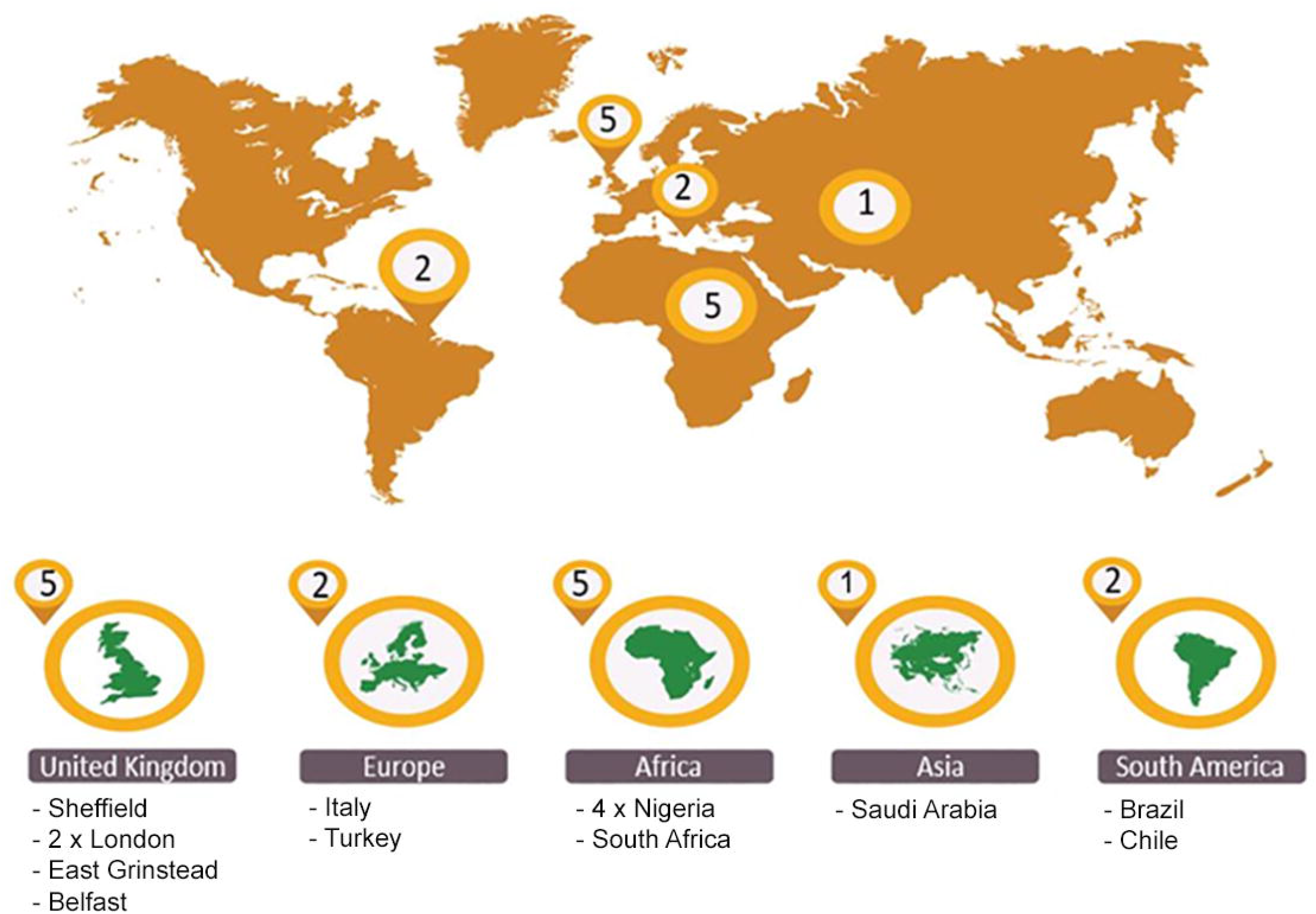
Geographical map of the centres participating in the study including five from the UK, one from Europe, five from Africa, two from Asia and two centres from South America.

All benign and malignant tumours in major and minor salivary glands were included, and clinical information about gender and age of the patient as well as the anatomical site of presentation (major gland or intra-oral location) was also obtained. Tumours involving the sinonasal region, nose and trachea were excluded. The data was anonymised locally before being shared for analysis. Cases with incomplete information were excluded after an initial review and duplicates and recurrences were also removed. Where possible, the histological diagnosis of the cases was reviewed (65% of cases). However due to the large number of cases in the cohort and the resource constraints, this was not possible for every case. The tumours were classified according to the 2017 WHO classification of salivary gland tumours (18). Twelve types of benign SGT were identified within the cohort, including common entities such as pleomorphic adenoma, Warthin tumour, basal cell adenoma and canalicular adenoma etc. For malignant SGT, 20 different tumour types were identified including mucoepidermoid carcinoma (MEC), adenoid cystic carcinoma (AdCC), acinic cell carcinoma (ACC), polymorphous adenocarcinoma (PAC), carcinoma ex pleomorphic adenoma (Ca ex PA) and adenocarcinoma NOS (AdNOS). Descriptive statistical analysis of the data was performed using frequencies and percentages of the variables in Microsoft Excel (2016). Student’s T test was used to determine statistical significance (where relevant).

## Results

### Overall

The total number of SGT was 5,798. Of these, 65% were benign tumours (n=3,751), and 35% (n=2,047) were malignant (Table). There was a slight female predilection (54%, n=3135) compared to 46% male patients (n=2,660). In both benign and malignant tumours, a higher incidence was noted in patients between the fourth to seventh decade of life accounting for 70% of the tumours (Figure 2). Most of the SGT involved the major glands 69% (n=3,966) with 31% (n=1,832) involving the minor glands. Parotid gland was the most common site of involvement (60%, n= 3,457). Within the minor glands, the palate was the most common location (60%, n= 1104) (Figure 3).

**Figure 2:**
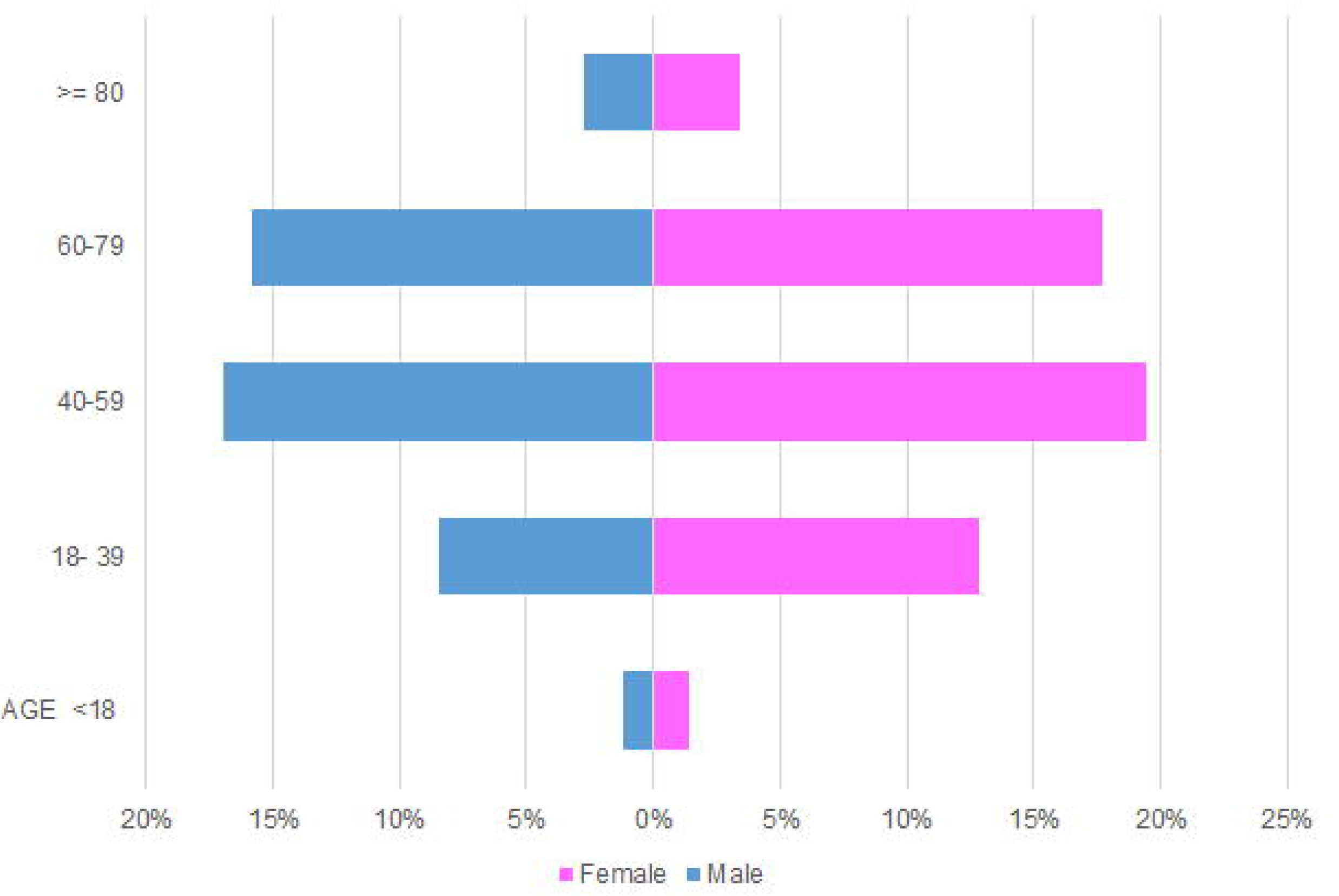
Age and gender distribution within the entire SGT cohort.

**Figure 3:**
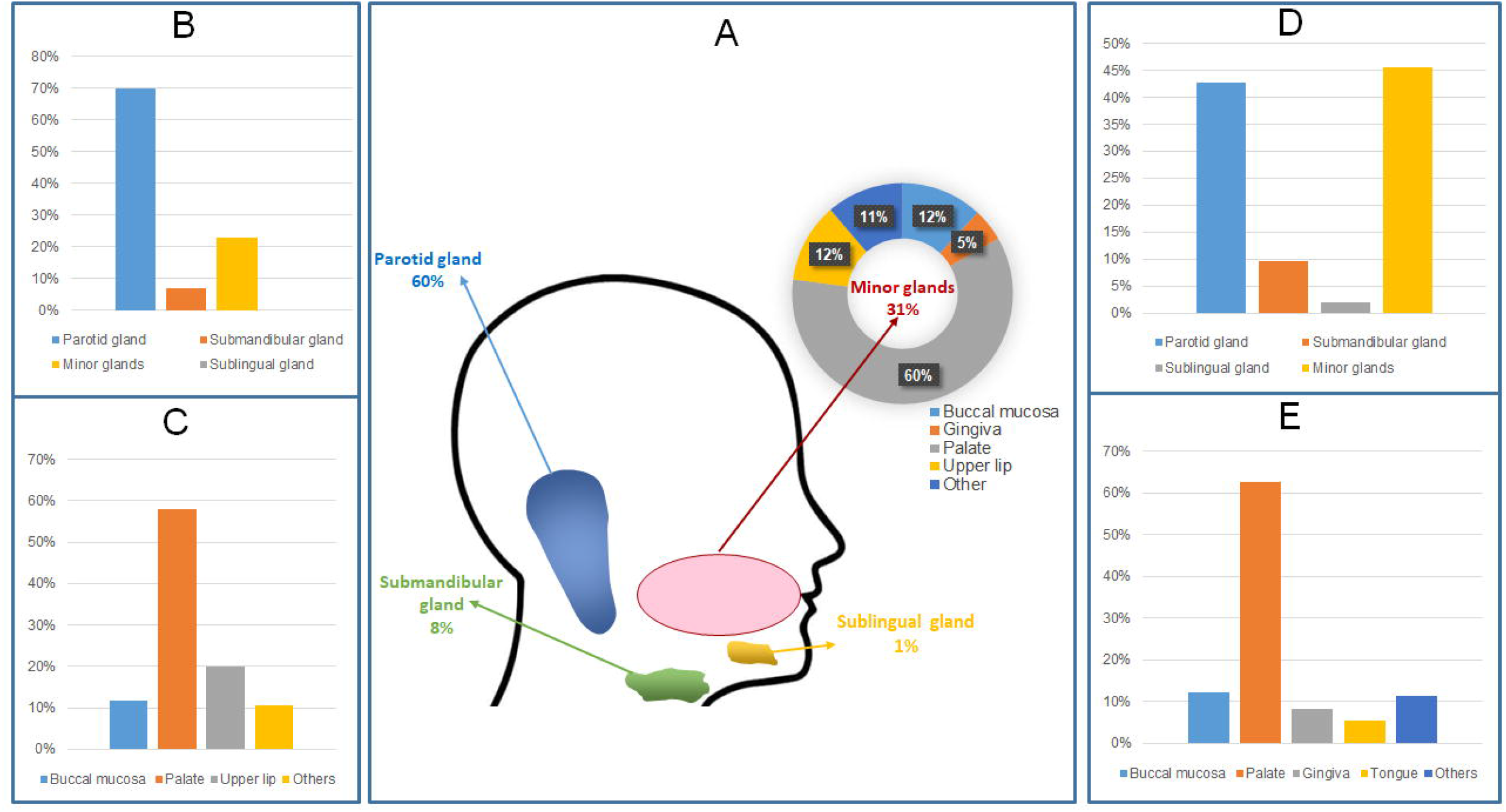
Anatomical site distribution of SGT. A) major and minor gland distribution across the entire cohort B) Benign tumours in all glands C) Benign tumours in minor glands D) Malignant tumours in all glands E) malignant tumours in minor glands.

**Table:** Benign and malignant SGT histological subtype distribution within the cohort.

| Diagnosis | Number | % of group | % of all tumours |
| --- | --- | --- | --- |
| **BENIGN** |  |  |  |
| Pleomorphic Adenoma | 2621 | 70% | 45% |
| Warthin Tumour | 623 | 17% | 11% |
| Basal Cell Adenoma | 172 | 5% | 3% |
| Canalicular Adenoma | 96 | 2% | 2% |
| Cystadenoma | 81 | 2% | 1% |
| Myoepithelioma | 62 | 2% | 1% |
| Oncocytoma | 43 | 1% | 1% |
| Other | 53 | 1% | 1% |
| **Benign total** | **3751** |  |  |
| **MALIGNANT** |  |  |  |
| Mucoepidermoid Carcinoma | 508 | 25% | 9% |
| Adenoid Cystic Carcinoma | 336 | 16% | 6% |
| Polymorphous Adenocarcinoma | 238 | 12% | 4% |
| Carcinoma ex pleomorphic adenoma | 215 | 11% | 4% |
| Acinic cell carcinoma | 185 | 9% | 3% |
| Adenocarcinoma NOS | 145 | 7% | 3% |
| Salivary duct carcinoma | 79 | 4% | 1% |
| MALT Lymphoma | 59 | 3% | 1% |
| Secretory Carcinoma | 51 | 2% | 1% |
| Epithelial Myoepithelial Carcinoma | 47 | 2% | 1% |
| Myoepithelial Carcinoma | 42 | 2% | 1% |
| Basal cell adenocarcinoma | 41 | 2% | 1% |
| Other | 101 | 5% | 2% |
| **Malignant total** | **2047** |  |  |

### Benign tumours

The most common benign tumour was pleomorphic adenoma accounting for 70% (n=2621) followed by Warthin tumour (17%, n=623), basal cell adenoma (3%, n= 172), canalicular adenoma (2%, n=96), cystadenoma (2%, n=81), myoepithelioma (2%, n=62) and oncocytoma (1%, n=43). All other rare benign tumours including sebaceous adenoma, ductal papilloma, sialoadenoma papilliferum and unclassified salivary tumours comprised 1% (n=53) of benign tumours (Table).

Comparison of the different geographical locations showed that pleomorphic adenoma was the most common benign tumour across all centres with a different range of incidence between centres ranging from 64% in Europe to 87% in South America. Warthin tumour was the second most common tumour for most of the centres with a range of incidence between 1% (African cohort) to 30% in the data from Europe. This was followed by basal cell adenoma accounting for 5% of the UK and 3% of both the European and African cases (Figure 4).

**Figure 4:**
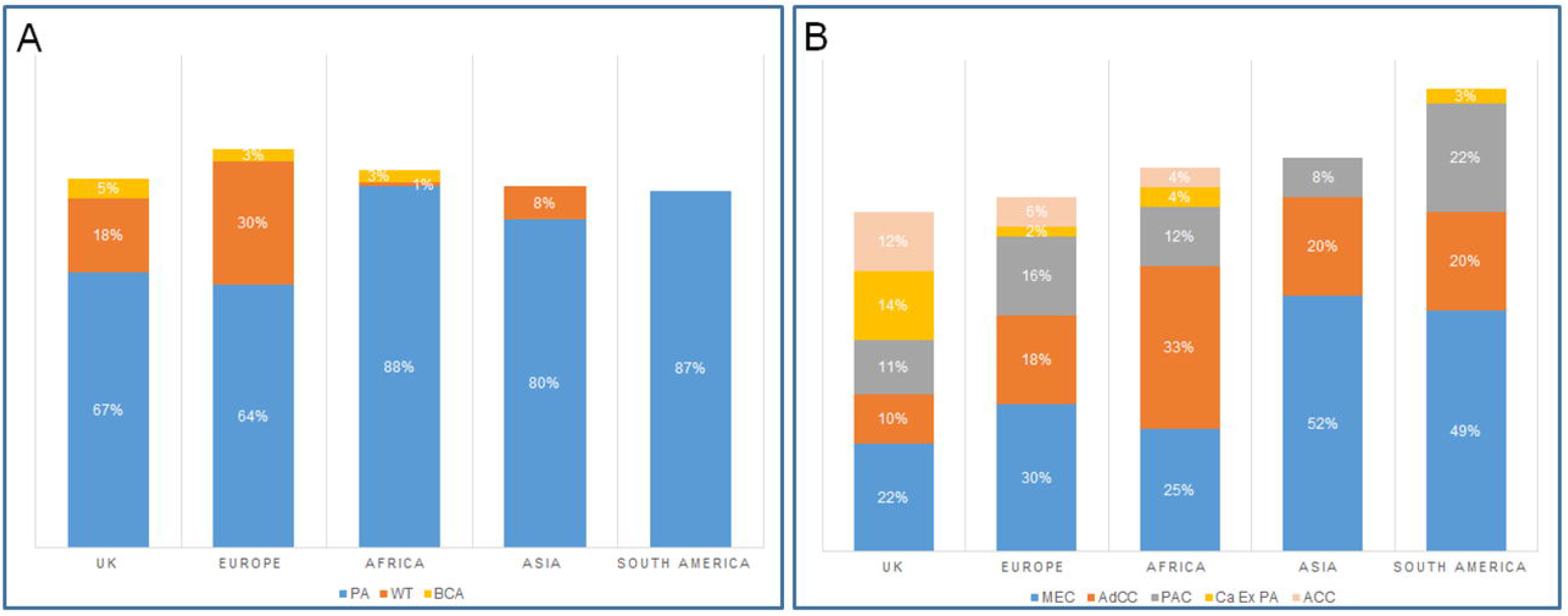
Geographic comparison of the most common benign and malignant SGT. A) benign tumours: PA-Pleomorphic adenoma, WT-Warthin tumour, BCA-Basal cell adenoma. B) malignant tumours: MEC- Mucoepidermoid carcinoma, AdCC- Adenoid cystic carcinoma, PAC- Polymorphous adenocarcinoma, Ca ex PA- Carcinoma ex pleomorphic adenoma, ACC- Acinic cell carcinoma.

### Gender distribution

For benign tumours, there was a slight female predilection (54%, n=2035) compared to males (46%, n=1716).

### Age distribution

The average age for benign tumours was 52 years with a female average age of 51.7 and male average of 52.4 years. The age range was very wide (1-98 years). The most commonly affected age group was 40-59 which accounted for 38% of benign tumours (n=1440) comprising 18% of male (n=692) and 20% female (n=748) patients respectively. The least affected age group was patients under 18 years of age who accounted for only 2% (n=82) of benign tumours (Figure 5).

**Figure 5:**
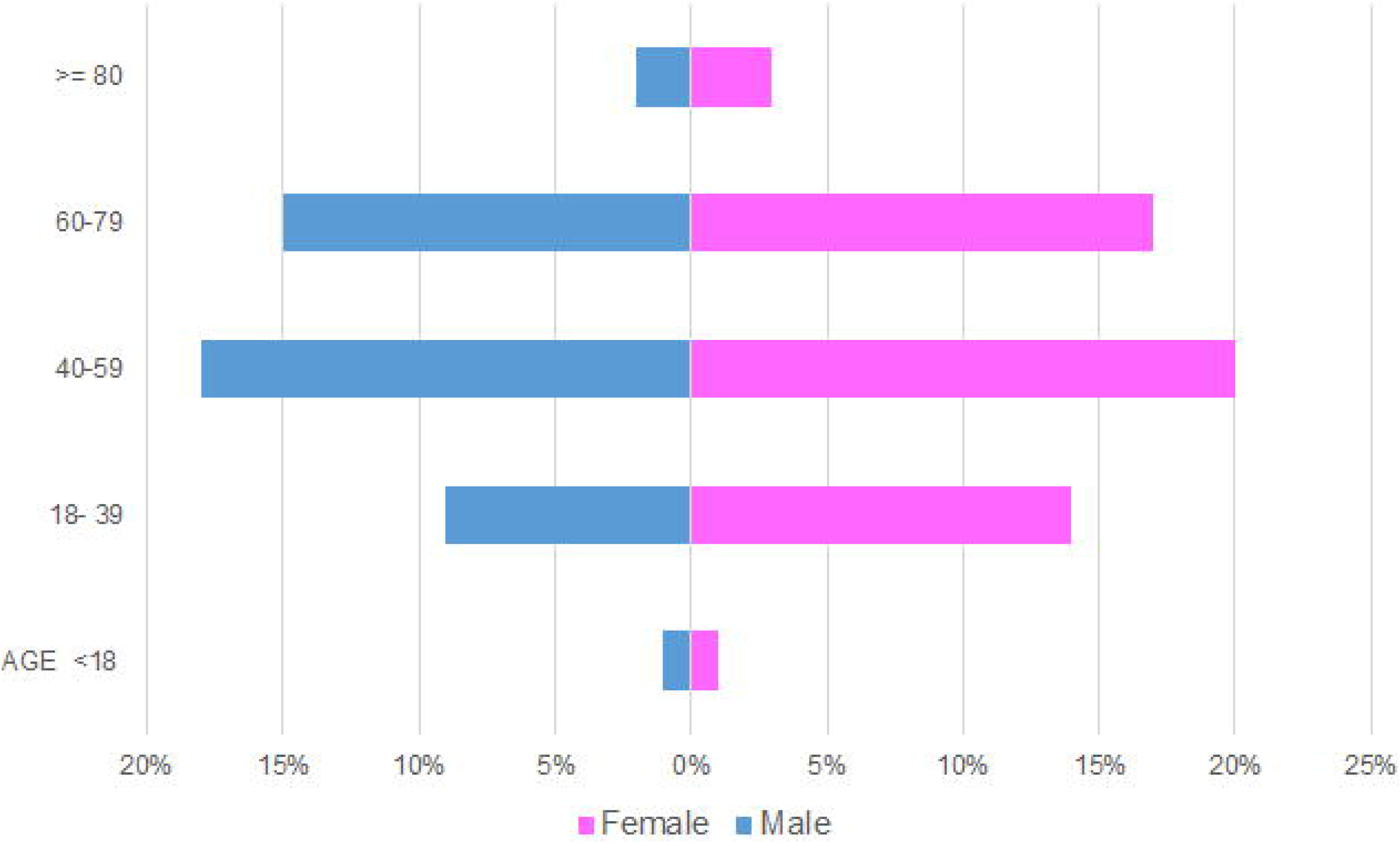
Age and gender distribution for benign salivary gland tumours.

### Site distribution

Benign tumours presented in both major and minor glands with the majority of tumours involving the parotid gland (70%, n= 2582), followed by minor glands (23%, n=899), submandibular gland (7%, n=265) and less than 1% (n=5) involving the sublingual gland. Within the minor glands, the palate was the most common location (58%, n=519). followed by upper lip (20%, n=178) and buccal mucosa (12%, n=104). Other sites included the floor of mouth, gingiva, lower lip, tongue and labial mucosa which altogether comprised 10% (n=98) of benign tumours (Figure 3).

The majority of pleomorphic adenomas involved the parotid gland (67%, n=1736), followed by minor glands (25%, n=645) and submandibular gland (8%, n=217). Only four cases (less than 1%) were identified in the sublingual gland. In the minor glands, (67%, n=433) of the pleomorphic adenomas involved the palate, followed by upper lip and buccal mucosa (12%, n=77, 11%, n=69) respectively. Warthin tumours were exclusive to the parotid gland (n=623). Most of the basal cell adenomas were reported in minor glands (55%, n=95), followed by parotid gland (40%, n=69) and submandibular gland (5%, n=8). In the minor glands, the upper lip was the most common minor gland site for basal cell adenomas (49%, n=47) (Figure 6).

**Figure 6.**
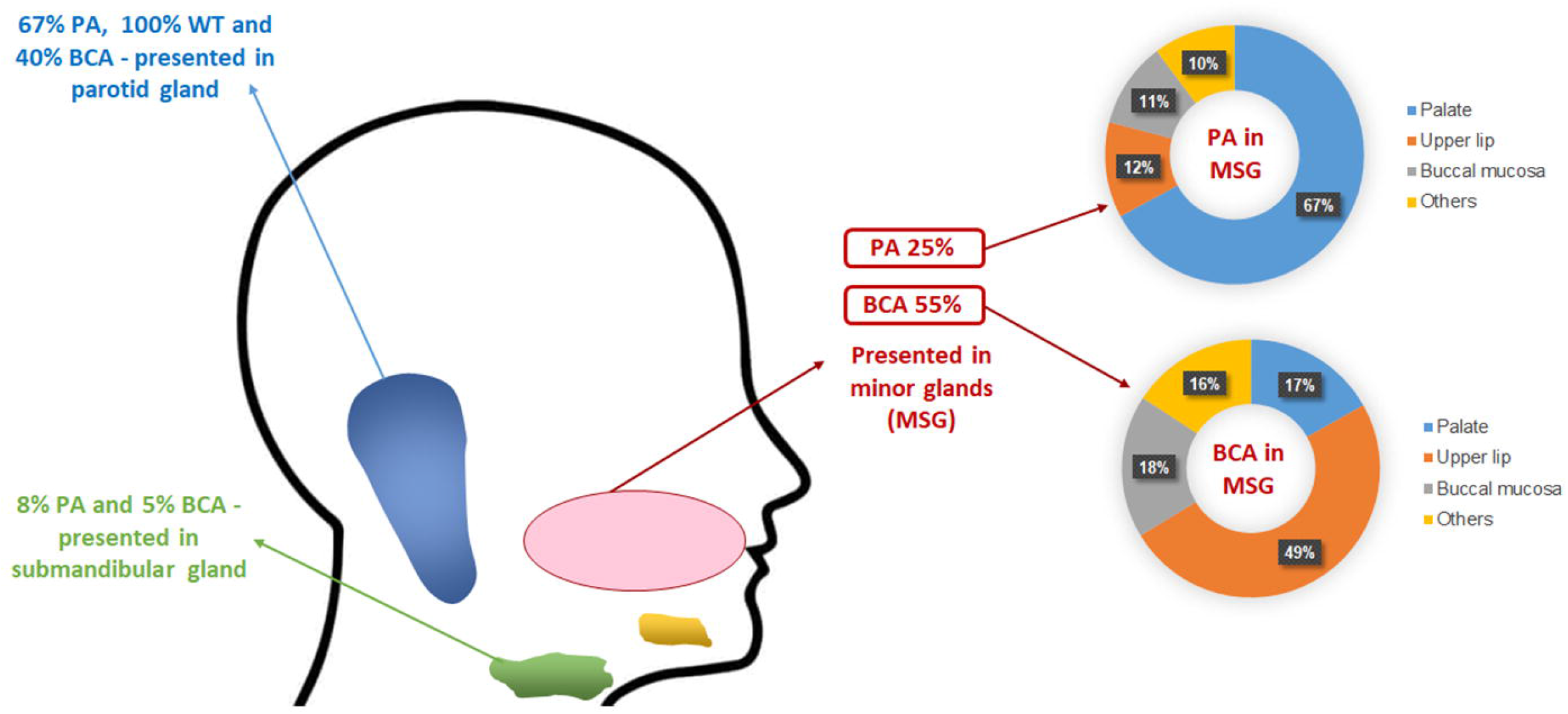
Anatomical site distribution of the most common benign SGT. Pleomorphic adenoma (PA), Warthin tumour (WT) and basal cell adenoma (BCA).

### Malignant tumours

The most common malignant tumour was mucoepidermoid carcinoma accounting for 25% of the malignant diagnoses (n= 508) followed by adenoid cystic carcinoma (16%, n=336), polymorphous adenocarcinoma (12%, n=238), carcinoma ex pleomorphic adenoma (11%, n=215), acinic cell carcinoma (9%, n=185), adenocarcinoma NOS (7%, n=145), salivary duct carcinoma (4%, n=79), MALT lymphoma (3%, n=59), secretory carcinoma (2%, n=51), epithelial myoepithelial carcinoma (2%, n=47), myoepithelial carcinoma (2%, n=42) and basal cell adenocarcinoma (2%, n=41). All other rare malignant tumours including, carcinosarcoma, clear cell carcinoma, cystadenocarcinoma, intraductal carcinoma, lymphoepithelial carcinoma, neuroendocrine carcinoma, oncocytic carcinoma, poorly differentiated carcinoma, sebaceous carcinoma, sialoblastoma and squamous cell carcinoma accounted for 5% of malignant tumours collectively (n=101) (Table).

Mucoepidermoid carcinoma was the most common malignant tumour in the majority of centres. The only exception was Africa, where adenoid cystic carcinoma was the most common tumour. Overall, adenoid cystic carcinoma was the second most common malignant SGT for most of the centres with a range of incidence of 10% in UK and 33% in African centres. (Figure 4).

### Gender distribution

There was a slight female predilection for malignant SGT (54% of malignant cases, n=1100) compared to males (46%, n=944).

### Age distribution

The average age for malignant tumours was 56 years with a female average of 54.4 and male average age of 57.4 years. As for benign tumours, the age range was very wide (1 to 108 years). The average age for malignant tumours was significantly higher than benign tumours (p<0.00001). The most affected age group was 60 to 79-year-old accounting for 36% (n=740) of malignant tumours, comprising 17% male (n=348) and 19% female (n=392) patients respectively. Similar to the benign tumours, the least affected age group was patients under 18 years of age (3%, n=69) (Figure 7).

**Figure 7:**
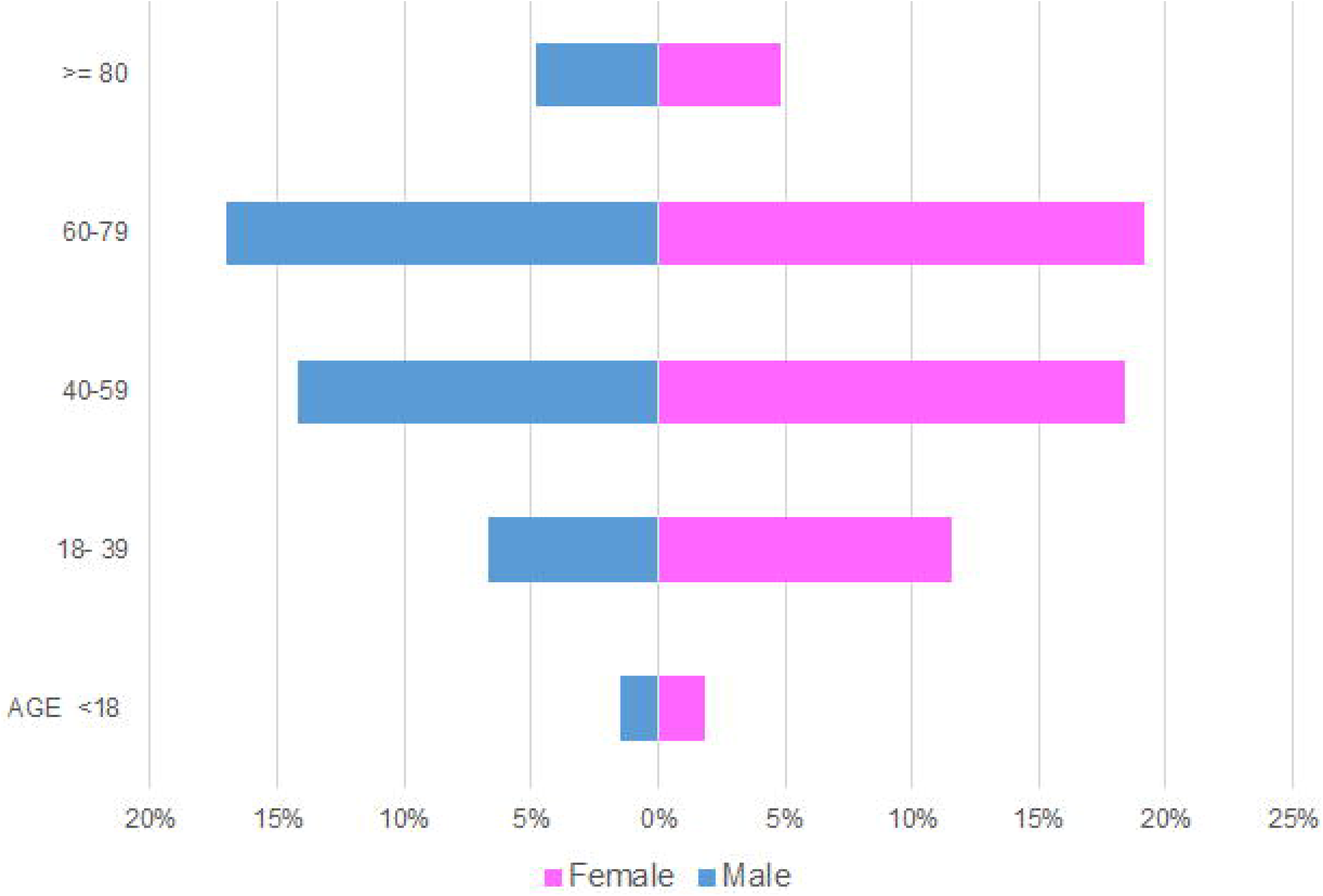
Age and gender distribution for malignant tumours.

### Site distribution

Minor glands were the most common site for malignant tumours (46%, n= 933), followed by parotid gland (43%, n=875), submandibular gland (10%, n=197) and sublingual gland (2%, n=42). Within the minor glands, the palate was the most common location (63%, n=585) followed by buccal mucosa (12%, n=114), gingiva (8%, n=78) and tongue (6%, n=51). Other rare sites included the floor of the mouth, labial mucosa, upper and lower lip, which accounted altogether for 11% of malignant diagnoses (n=105) (Figure 3).

The majority of mucoepidermoid carcinomas involved the minor glands (56%, n=285), followed by the parotid gland (34%, n=173), submandibular glands (7%, n=36) and sublingual glands (3%, n=12). In the minor glands, 60% (n=172) of the mucoepidermoid carcinomas occurred in the palate. Most of the adenoid cystic carcinomas (60%, n=203) involved the minor glands, followed by parotid (26%, n=87), submandibular (10%, n=33) and sublingual (4%, n=13) glands. Within the minor glands, 70% of adenoid cystic carcinomas presented in the palate (n=141). Most of the polymorphous adenocarcinomas presented in the parotid gland (47%, n=111), followed by the minor (46%, n=110) and submandibular (7%, n=17) glands. In the minor glands, the palate was the most common site for the polymorphous adenocarcinomas accounting for (72%, n=80). The majority of carcinoma ex pleomorphic adenomas involved the minor glands (61%, n=132), followed by the parotid gland (32%, n=70), submandibular gland (7%, n=13) and only one case was reported in the sublingual gland. In the minor glands, 62% (n=82) of carcinoma ex pleomorphic adenomas occurred in the palate. Parotid gland was also the most common site for acinic cell carcinoma (65%, n=120), followed by the minor glands (23%, n=42), submandibular gland (9%, n=17) and sublingual gland (3%, n=5). The palate was the most common minor gland site for acinic cell carcinomas (50%, n=21) (Figure 8).

**Figure 8:**
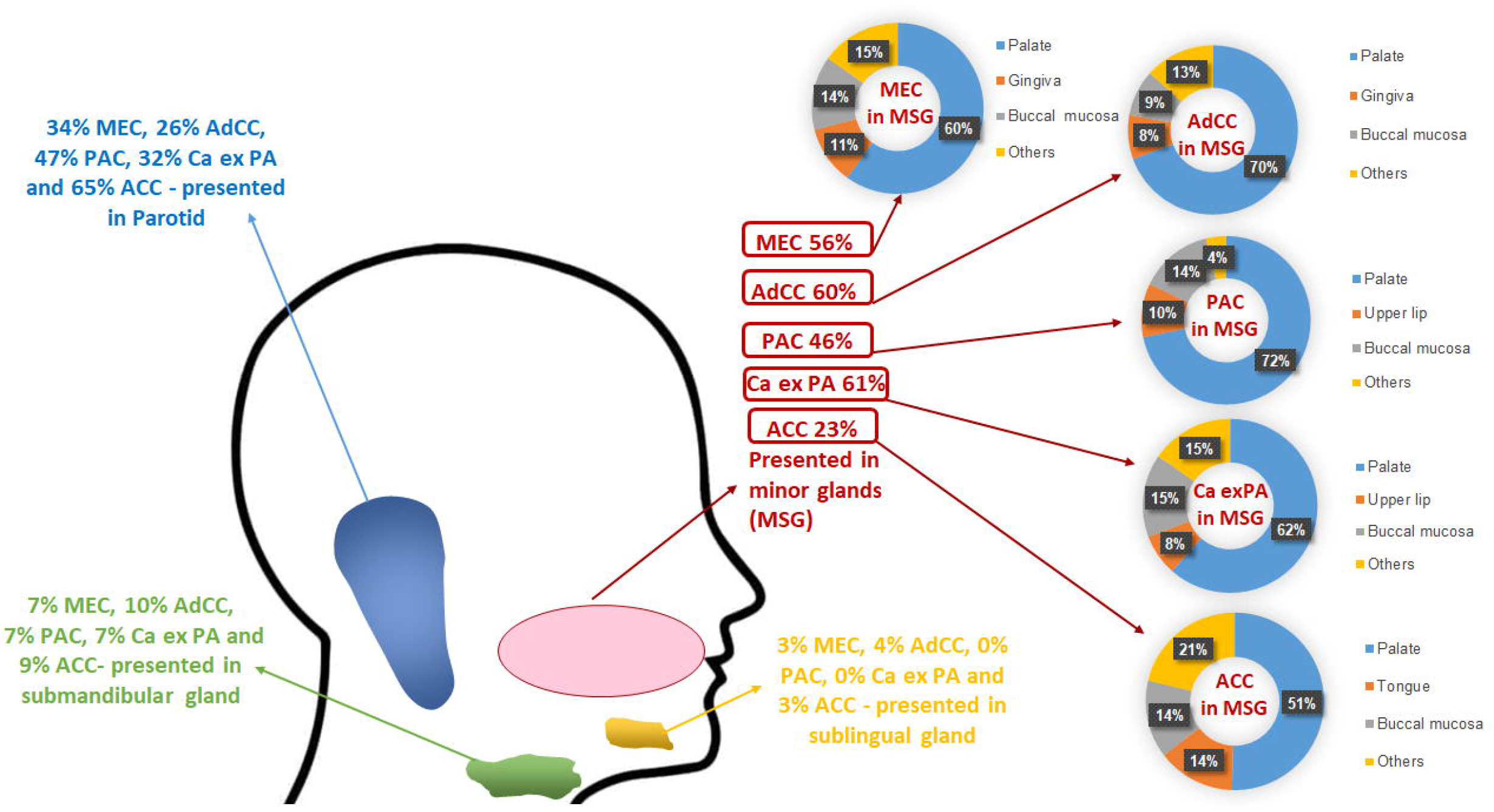
Anatomical site distribution of the most common malignant SGT. Mucoepidermoid carcinoma (MEC), adenoid cystic carcinoma (AdCC), polymorphous adenocarcinoma (PAC), carcinoma ex pleomorphic adenoma (Ca ex PA) and acinic cell carcinoma (ACC).

## Discussion

SGT are rare and our study is one of the most extensive recent reports investigating the incidence and demographics of these tumours from multiple centres and the majority of the WHO geographical regions. We collected 5798 cases from pathology centres across the world and found that benign tumours accounted for the majority of SGT (65%) compared to malignant tumours (35%). This finding is consistent with the existing literature [10,14,19–21]. There was a slight predilection for females in our study, which has also been reported by some other studies [4,17,21–22]. However, some single centres studies have reported a higher incidence of salivary gland tumours in males or an equal gender involvement [9,10,12–13]. Furthermore, we found a peak incidence of SGT between the fourth to seventh decades of life (approximately 70% of cases) which is similar to findings previously reported in the literature [4,10,17,22].

Overall, the parotid gland was the most common location for SGTs accounting for (60%) followed by minor (31%) and submandibular (8%) salivary glands similar to older and similarly large SGT demographical studies [10,17,19–23]. Our study shows low prevalence of sublingual tumours (47 out of 5623), similar to findings reported by Satko et al., 2000 and Eveson and Cawson (1985) [20,23].

### Benign SGT

Our results show that pleomorphic adenoma was the most common benign tumour; this is consistent with prevalence rates reported in the literature [4,5,8,10,12,14,17, 21]. Comparison of incidence between centres showed pleomorphic adenoma to be the most common benign tumour in all geographical locations included in the study ranging from 64% in Europe to 87% in South America. Warthin tumour was the second most common benign tumour which is also similar to earlier demographical studies [8,11,17,22,23,25]. The present study shows that basal cell adenoma was the third most common benign tumour; however accurate comparisons between previous studies of basal cell and canalicular adenomas is difficult as older studies tended to combine these into a single group of monomorphic adenomas [7,21,25]. Curiously no basal cell adenomas were reported in Asia and South America, whereas they comprised 5% of benign SGT in UK and 3% in both European and African centres.

In the literature, benign tumours have been reported to occur more in younger and female patients [22]. Our study shows a similar finding where benign tumours appeared to be more common in female patients; however, the average ages were quite similar (51.7 years in female compared to 52.4 in male patients).

The majority of the benign tumours were located in the parotid gland (70%), followed by minor [23%] and submandibular (7%) glands. We found only five benign cases (less than 1%) involving the sublingual gland. Similar findings were seen in other older SGT studies with some variations in distribution across different anatomical sites [4,10,14,17,21–22].

The most common location for the pleomorphic adenoma was the parotid gland (67%), followed by minor glands and the submandibular gland (25% and 8% respectively. This observation was consistent with previous reports [4,11,13,17,22]. Warthin tumour was seen exclusively in the parotid gland (100%).This observation is similar to other studies suggesting that this tumour exclusively involves parotid gland [11,13,22]. Basal cell adenomas were reported in minor glands in 55% of the cases and about 40% were seen in the parotid gland. Accurate comparison with the literature was not possible due to the small number of reported studies and cases.

### Malignant SGT

Mucoepidermoid carcinoma was the most common malignant salivary gland tumour accounting for 25% of diagnoses; this is in agreement with previously reported prevalence rates worldwide [4,8,21,22]. We found that mucoepidermoid carcinoma was the most common tumour in all centres except Africa where adenoid cystic carcinoma (33%) was the most common malignant diagnoses. Bello *et al*., 2012, reported somewhat similar findings with adenoid cystic carcinoma as the most common malignant tumour in their 2012 study [17]. However, this was limited to data from two centres and did not examine the geographical differences in depth. The second most common malignant diagnosis for most of the centres in our study was adenoid cystic carcinoma accounting for 16% of all malignancies similar to other reports [5,8,11–12,21–24] although some variations in its incidence have been reported. The next most common diagnoses were polymorphous adenocarcinoma and carcinoma ex pleomorphic adenoma accounting for 12 and 11% of diagnoses respectively. In fact, there is significant variability in the literature about the incidence of these two entities which is perhaps related to small and unicentric cohorts. Similarly, acinic cell carcinoma was the fifth most common diagnosis (9%) although, some studies have reported it as the third most common malignant SGT [4,5] however these findings were also based on a small cohort size from only one centre. Adenocarcinoma NOS was the next most common diagnosis accounting for 7% of cases which may not reflect the exact incidence of this entity as development of ancillary molecular and sequencing techniques has led to more specific diagnoses and reduction in the use of this diagnosis [26].

Malignant SGT are more likely to occur in older patients [22–23]. Our study shows similar results where the average age for malignant tumours was significantly higher than benign tumours (p<0.00001). Malignant SGT involved the parotid gland in about 43% of cases, which aligns with known literature [11,14]. The incidence of malignant tumours in minor salivary glands was slightly higher than the parotid gland (i.e. 46%), highlighting the importance of considering malignant SGT in differential diagnoses [14,28]. In the present study, 42 out of 47 cases involving the sublingual gland were malignant tumours, which is similar to Tian et al.’s findings from 2010 where they reported 95% of sublingual tumours to be malignant [10].

The most common location for mucoepidermoid carcinoma was minor glands (56%), followed by parotid, submandibular and sublingual (34%, 7% and 3% respectively) glands which is in agreement with the findings of Jones et al., 2008 but in disagreement with other reports which have found parotid to be the most common site [11,13,32]. This could be due to the fact that a number of these cases were reported in specialist oral and maxillofacial pathology units (some of which usually only report intra-oral specimens) although many of the included centres also have expertise in head and neck (H&N) pathology.

Understanding the epidemiological landscape and distribution of histological subtypes of salivary gland tumours is crucial for a better diagnosis of this diverse and complex group of tumours. It would be useful for future studies to include more geographical locations as well as other H&N and ENT centres. It would also be useful to establish international datasets of these tumours for use by other researchers (whole slide image or research repositories) including histological reassessment and classification according to the most recent WHO criteria.

## Conclusions

Salivary gland tumours are rare, but show a gradual increasing incidence over the last decade and a half. We report the largest multicentre investigation of SGT to date showing that the majority are benign (65%), with a slight predilection for females (54%). Approximately 70% of SGT occur in patients between the fourth to seventh decade of life with a significant difference between the average age for benign and malignant tumours. Pleomorphic adenoma was the most common benign and mucoepidermoid carcinoma the most common malignant tumour. The majority of SGT presented in the major glands (69%), with the parotid gland being the most common location (70%)for benign and minor glands (46%) for malignant tumours. More extensive studies of SGT need to be conducted to understand and update the epidemiological landscape of these tumours and correlate it with prognosis.

## DECLARATIONS

### Availability of data and material

Anonymised data and materials available upon request and through the publication.

### Code availability

Not applicable

### Authors’ contributions

Study design (IA, SAK), Data collection and analysis (all authors), Manuscript preparation (IA, SAK), Manuscript review (all authors).

Additional declarations for articles in life science journals that report the results of studies involving humans and/or animals - Not applicable

### Ethics approval

In place. Reference- 20/WS/0017

### Consent to participate

Not applicable

### Consent for publication

Not applicable

